# Bioinformatic evaluation of the potential oral-gut translocation of periodontal pathogens in patients with colorectal polyps

**DOI:** 10.1101/2024.04.29.591540

**Authors:** Naoki Takahashi, Marin Yamaguchi, Keisuke Sato, Takahiro Tsuzuno, Shuhei Mineo, Nao Nakajima, Kazuya Takahashi, Hiroki Sato, Haruna Miyazawa, Yukari Aoki-Nonaka, Yutaro Ito, Koji Taniguchi, Shuji Terai, Kohei Ito, Koichi Tabeta

**Author notes:** Both the authors have contributed equally to the work (Naoki Takahashi, Marin Yamaguchi). Corresponding author: Naoki Takahashi, Division of Periodontology, Niigata University Graduate School of Medical and Dental Sciences, Niigata, 951-8514, Japan, Kohei Ito, BIOTA Inc., Tokyo, 101-0022, Japan.

## Abstract

**Objective:** This study aimed to characterize the profiles of the oral and gut microbiota of patients with colorectal polyps using 16S rRNA gene sequencing and bioinformatic approaches.

**Background:** Previous studies have shown microbial translocation from the oral cavity to the gut, implying pathogenic impacts on gastroesophageal disease, including colorectal cancer (CRC). However, its details remain unclear.

**Methods:** Twenty patients scheduled for endoscopic colorectal polypectomy were enrolled in this study. Oral samples (saliva and subgingival dental plaque) and intestinal samples (feces and swab of intestinal mucosa) were collected during preoperative and 6-month-postoperative reassessment periods. After sequencing the V3–V4 region of the bacterial 16S rRNA gene, several bioinformatic analyses (bacterial composition, diversity, core microbiome, and shared ASV) were performed on pre– and postoperative samples for each subject.

**Results:** The bacterial composition was dominated by *Bacteroides*, *Streptococcus*, *Fusobacterium*, *Veillonella*, and *Prevotella_7* in all four samples. Beta diversity analysis using weighted UniFrac distance distinctly segregated the samples between oral and intestinal environments in the principal coordinate analysis plot. Core microbiome analysis revealed that *Streptococcus* and *Porphyromonas* were dominantly shared in intra-oral environments. Additionally, alongside *Streptococcus*, periodontitis-related bacteria, such as *Veillonella*, *Fusobacterium*, *Porphyromonas*, *Prevotella_7*, *Haemophilus*, and *Prevotella*, were identified as shared genera between oral and intestinal environments. Finally, shared ASV analysis demonstrated that *Streptococcus* was shared in the oral and intestinal environments of most patients, while periodontal pathogens were shared in some patients.

**Conclusions:** The core microbiome and shared ASV analyses revealed that several genes are shared between oral and intestinal environments in patients with colorectal polyps, indicating the oral–gut translocation of periodontitis-related bacteria. Further large-scale studies are needed to elucidate their involvement in CRC.

## Introduction

Periodontitis, a chronic inflammatory disease affecting the tissue around teeth, can lead to alveolar bone destruction and eventual tooth loss if untreated^1^. A recent systematic review found that the global prevalence of periodontitis has increased to approximately 60% in the last decade (2011–2020), surpassing previous estimates from 1990 to 2010^2^. Its adverse effects extend beyond the oral cavity, influencing the onset and progression of systemic diseases such as cardiovascular disease, respiratory infections, and diabetes^3,4^. Emerging research on the oral-gut microbial and immune axis suggests a link between periodontitis and gastrointestinal inflammatory diseases and cancer^5^. Specific bacterial infections, including pathogenic *Escherichia coli*, *Salmonella enterica*, *Helicobacter pylori*, and *Fusobacterium nucleatum*, have been associated with colorectal cancer (CRC) risk^6^. *F. nucleatum*, an oral commensal periodontal pathogen, is frequently found in CRC tissues and is linked to tumor progression and prognosis by promoting tumor growth and metastasis^7,8^. High-resolution microbiome analysis suggests extensive translocation of oral bacteria to the intestine^9^. The oral-gut translocation of periodontitis-related pathogens, such as *Porphyromonas*, *Prevotella*, *Neisseria*, *Veillonella*, and *Haemophilus*, has also been reported^10^. Additionally, ectopic gut colonization of oral bacteria like Klebsiella and Enterobacter species exacerbates intestinal diseases^11,12^. Although most gut microbiota studies in humans use fecal specimens, the luminal and mucosal microbiota in the intestinal tract exhibit distinct compositions^13–15^. Intestinal mucosa-associated microbiota (MAM) interacts more directly with host intestinal epithelial cells compared to luminal-associated microbiota (LAM), suggesting MAM provides more precise information in predicting bacteria associated with intestinal diseases^16–18^. However, research on the association of periodontal pathogens with MAM in colorectal polyps is limited. This study aims to investigate the involvement of periodontopathogens in CRC by evaluating oral (saliva and subgingival dental plaque) and intestinal (feces and intestinal mucosa swabs) samples in Japanese patients with colorectal polyps using bioinformatic approaches.

## Materials and methods

### Study Design

This study comprised 20 patients who underwent endoscopic mucosal resection of colorectal lesions at the Division of Gastroenterology and Hepatology, Graduate School of Medical and Dental Sciences, Niigata University. Oral (saliva and subgingival dental plaque) and intestinal samples (feces and swab of intestinal mucosa) were collected before the operation and during reassessment six months post-surgery. Patients who used antibiotics within the past three months were excluded. Saliva collection involved patients spitting into a sterile Falcon tube for 5 min. Subgingival dental plaque samples were collected using two sterile paper points inserted into the gingival sulcus for 10 s. The preoperative MAM was obtained by swabbing the surface of the polyp before excision. Postoperative sampling involved swabbing the intestinal mucosa from which the polyp was excised. All samples were promptly stored at −80°C after collection. Some postoperative samples could not be collected due to unexpected hospital transfers and/or changes in the patient’s treatment plan, resulting in postoperative analysis being performed with 13 samples for saliva, subgingival oral plaque, and feces, and ten samples for intestinal mucosal tissues. Patient characteristics are shown in **Table 1**.

**Table 1.** Patients characteristics.

|  |  |
| --- | --- |
| Remaining Teeth | $21.95 \pm 8.04$ |
| Plaque Control Record | $46.08 \pm 25.07$ |
| Mean Probing depth | $2.935 \pm 0.48$ |
| Bleeding On Probing | $15.6 \pm 14.8$ |
| Deepest Probing depth | $5.75 \pm 1.59$ |
| Colorectal Neoplasm Macroscopy |  |
| Flat | 13 |
| Raised | 17 |
| Location |  |
| Right | 10 |
| Left | 10 |
| Cancer or adenoma |  |
| Cancer | 14 |
| Adenoma | 6 |

### Total DNA extraction and high-throughput sequencing

Samples suspended in preservation solution were homogenized, then the supernatant was boiled for 5 min. Total DNA was extracted using Genefind V2 (Beckman Coulter, United States) according to the manufacturer’s protocol. PCR reactions were conducted with the universal bacterial primers 341F-806R (F5′-TCGTCGGCAGCGTCAGATGTGTATAAGAGACAGNNCCTACGGGNGGCW GCAG-3′) and (R 5′-GTCTCGTGGGCTCGGAGATGTGTATAAGAGACAGNNGACTACHVGGGTAT CTAATCC-3′), to amplify the V3-V4 of the 16S rRNA gene. The thermal conditions were 95°C for 3 min, followed by 95°C for 30 s, 55°C for 30 s, and 72°C for 30 s, with a final extension at 72°C for 5 min. Purification was carried out using AmpureXP (Beckman Coulter), and the primers were removed. The second set of PCR reactions were conducted with the primers (F5′-AATGATACGGCGACCACCGAGATCTACAC TATAGCCT TCGTCGGCAGCGTC-3′ and R5′-CAAGCAGAAGACGGCATACGAGAT CTAGTACG GTCTCGTGGGCTCGG-3′) to attach the necessary adapter sequences and Unique Dual Index for library preparation. The thermal conditions were similar to the first set of PCR reactions. After purification of the PCR products, quantification was carried out using gel electrophoresis and Qubit (Invitrogen, Waltham, MA, USA). Adjustments were made to ensure equal concentration among all samples. DNA samples, library preparation, and amplicon sequencing were performed using 301 bp × 2 pair-end sequencing on MiSeq Reagent Kit V3 (Illumina Inc., San Diego, CA, USA) and the Illumina MiSeq platform (Illumina Inc.) at Genome-Lead Co., Ltd. (Kagawa, Japan).

### Microbiome analysis

Microbiome analysis was performed based on a previous study^19^. Briefly, raw FASTQ data were imported into the QIIME2 platform (version 2023.5) as qza files^20^. Denoising sequences and quality control were performed using the QIIME dada2. Finally, the sequences were produced into amplicon sequence variants (ASVs)^21^. ASVs were assigned to the SILVA database’s SSU 138 using the QIIME feature-classifier classification scikit-learn package^22^. The subsequent analysis excluded ASVs classified as mitochondria, chloroplast, or unassigned. The alpha diversity index Shannon diversity indices were calculated using the diversity plugin. The beta diversity index weighted and unweighted UniFrac distances were calculated, and the microbial community structure differences between the groups were visualized following a principal coordinate analysis (PCoA). The data were visualized using ggplot2 (version 3.4.4) ^23^, ggprism (version 1.0.4) (https://csdaw.github.io/ggprism/), matplotlib (version 3.7.1)^24^, and seaborn (version 0.12.2)^25^.

### Calculation of shared ASVs

Shared ASVs analysis was performed based on a previous study^26,27^. We defined shared ASVs as ASVs shared by >1.0% of both samples in this study. The calculation was conducted using the custom Python code (q2-shared_asv, https://github.com/biota-inc/q2-shared_asv) with 0.01 for p-percentage.

### Statistical analysis

Mann–Whitney U test was used to compare alpha diversity (Shannon diversity index) and pairwise UniFrac distance. For comparing differences in beta diversity (weighted UniFrac distance) between samples, all PERMANOVA analyses were conducted with 5,000 trials to assess the statistical significance. Multiple testing corrections were applied by computing False Discovery Rates (FDR) using the Benjamini–Hochberg method, with *Q*-values (adjusted P-values) <0.05 considered statistically significant. The core microbiome threshold was defined as a ratio of 10%^28^. In this experiment, the core microbiome was identified based on the ratio of total counts to all subject pairs for each comparison >10.0% and at a relative abundance of >1.0% at the genus level. For shared ASV analysis, a similar threshold was defined with a relative abundance of at least 1.0%. Differentially abundant microbial taxa were identified using the Analysis of Composition of Microbiomes (ANCOM) method in QIIME 2. The *W*-statistic indicates the strength of the ANCOM test for the tested number of genera^29^. Hierarchical clustering was performed using Euclidean distance and Ward’s method.

## Results

### Major bacterial taxonomic composition of gut and oral samples

We initially analyzed the overall relative abundance of bacteria and their taxonomic composition at the genus level across the four samples: saliva, dental plaque, feces, and swab of intestinal mucosa (**Fig. 1**). This analysis was conducted on pre– and postoperative samples for each subject. The genera *Bacteroides*, *Streptococcus*, *Fusobacterium*, *Veillonella*, and *Prevotella_7* were the most abundant across all four samples. Additionally, we examined changes in bacterial abundance between pre– and postoperative samples for each using ANCOM **(Fig. S1**). However, no periodontal pathogens with elevated w-statistics were observed in either oral or intestinal samples. These results suggest that bacterial abundance remains mostly consistent between pre– and postoperative periods.

**Figure 1.**
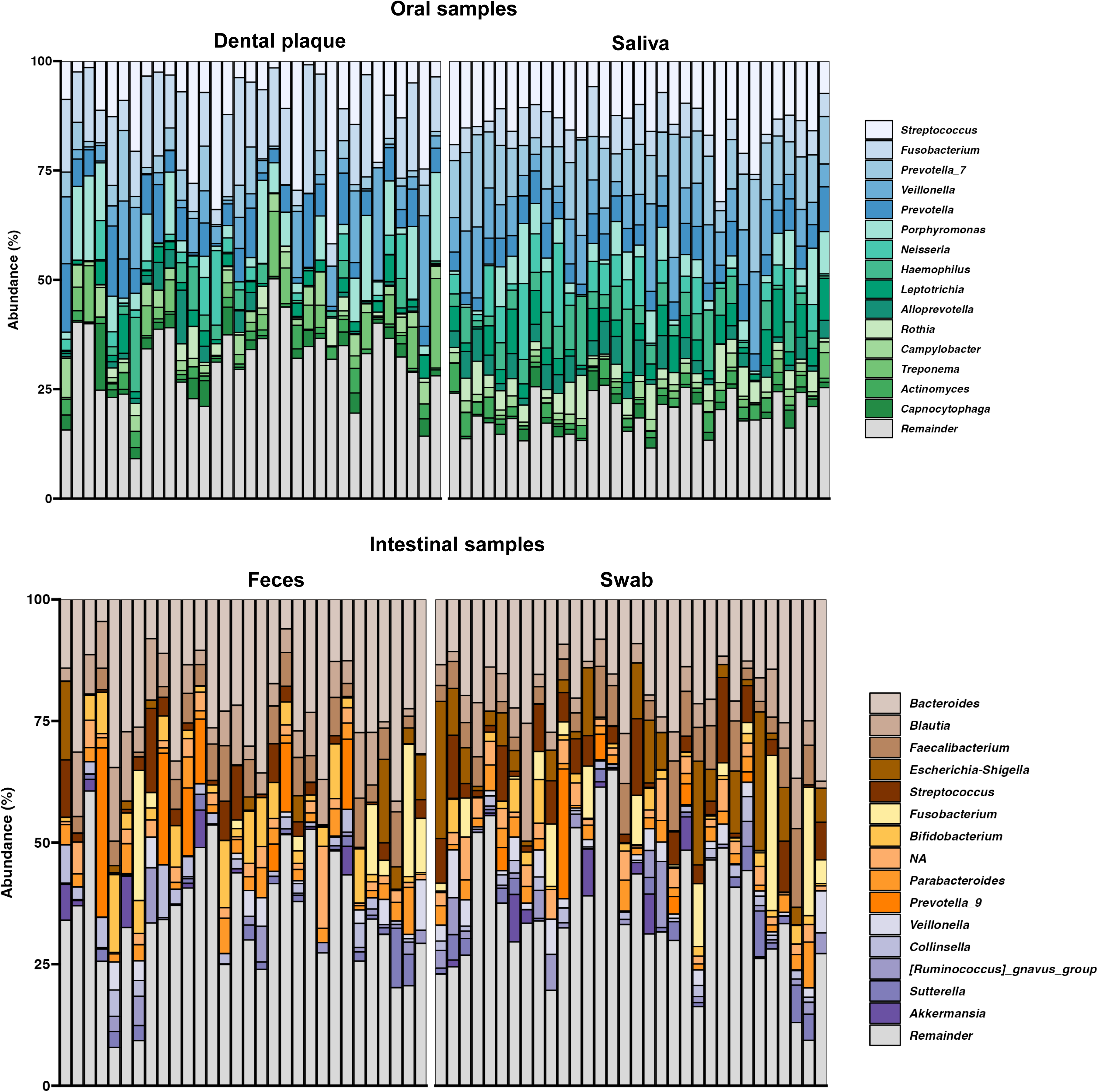
Bacterial composition of each sample. Bacterial composition in saliva, subgingival dental plaque, feces, and swab of the intestinal mucosa of Japanese patients with colorectal neoplastic lesions (n = 33 for oral bacteria, saliva, swab; n = 30 for feces).

### The bacterial diversity of samples was segregated between oral and intestinal environments

Bacterial diversity analysis was done between pre– and post-operation for each sample. For the alpha diversity analysis, the Shannon diversity index showed no significant changes, as indicated by the similarities in violin plots with pre– and postoperative sampling time as the x-axis and the Shannon diversity index as the y-axis for all sample locations (**Fig. 2A**). For the beta diversity analysis, the PCoA plot using unweighted UniFrac distance distinctly segregated the samples between oral and intestinal environments (**Fig. 2B**).

**Figure 2.**
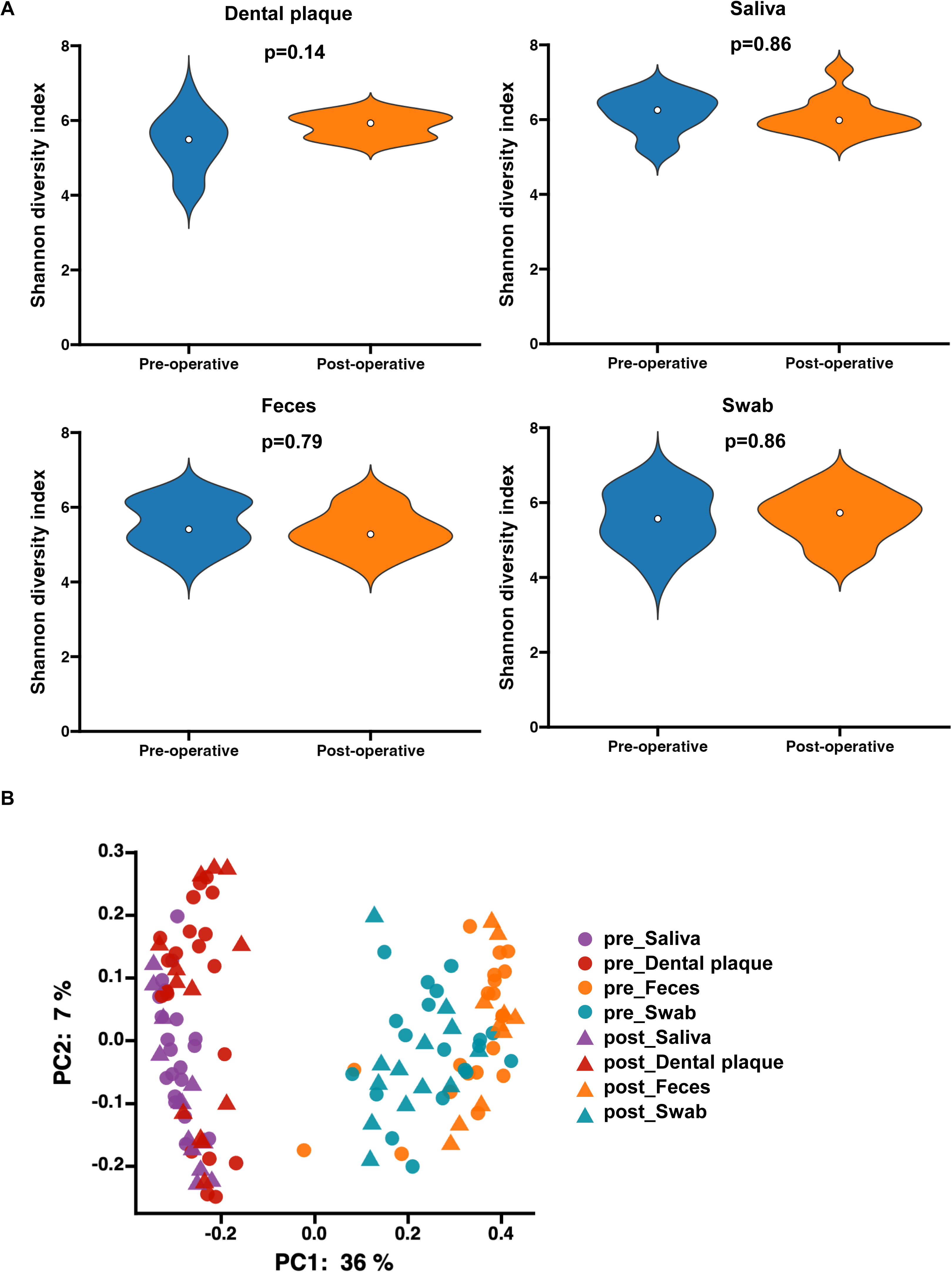
Diversity of the microbiome of each sample. Alpha and beta diversity of the microbiome of each sample. (A) Shannon diversity indices. (B) PCoA plot of microbiomes of each sample based on weighted UniFrac distance. P-values were calculated by testing the pre– and postoperative values within each sample and correcting for multiplicity.

### Some periodontal pathogens were detected between gut and oral samples as core taxa

Expanding on insights from microbial compositions of oral and intestinal samples, we conducted core microbiome analysis to identify intra-individual sharing of microbial taxa at the genus level. This analysis encompassed not only comparisons between the same samples pre– and post-operation but also between different samples pre– and post-operation.

Total counts were calculated by summing the number of patients where the same genus had a presence of at least 1% in intra-individual comparisons (**Fig. 3A**, **Supplementary Table. S1**). Ratios of total counts to all patient pairs for each comparison were then calculated (**Supplementary Table. 1**, **2**), and the top 15 genera exceeding the threshold were extracted as core microbiome (See Materials and Methods). These top 15 genera include *Streptococcus*, *Bacteroides*, *Veillonella*, *Fusobacterium*, *Faecalibacterium*, *Blautia*, *Porphyromonas*, *Prevotella_7*, *Haemophilus*, *Prevotella*, *Neisseria*, *Parabacteroides*, *Prevotella_9*, *Collinsella*, and *Bifidobacterium*. Notably, *Veillonella*, *Fusobacterium*, *Piorphyromonas*, *Prevotella_*7, *Haemophilus*, and *Prevotella* are specifically defined as periodontal pathogens, demonstrating their core microbiome presence in any of the comparisons.

**Figure 3.**
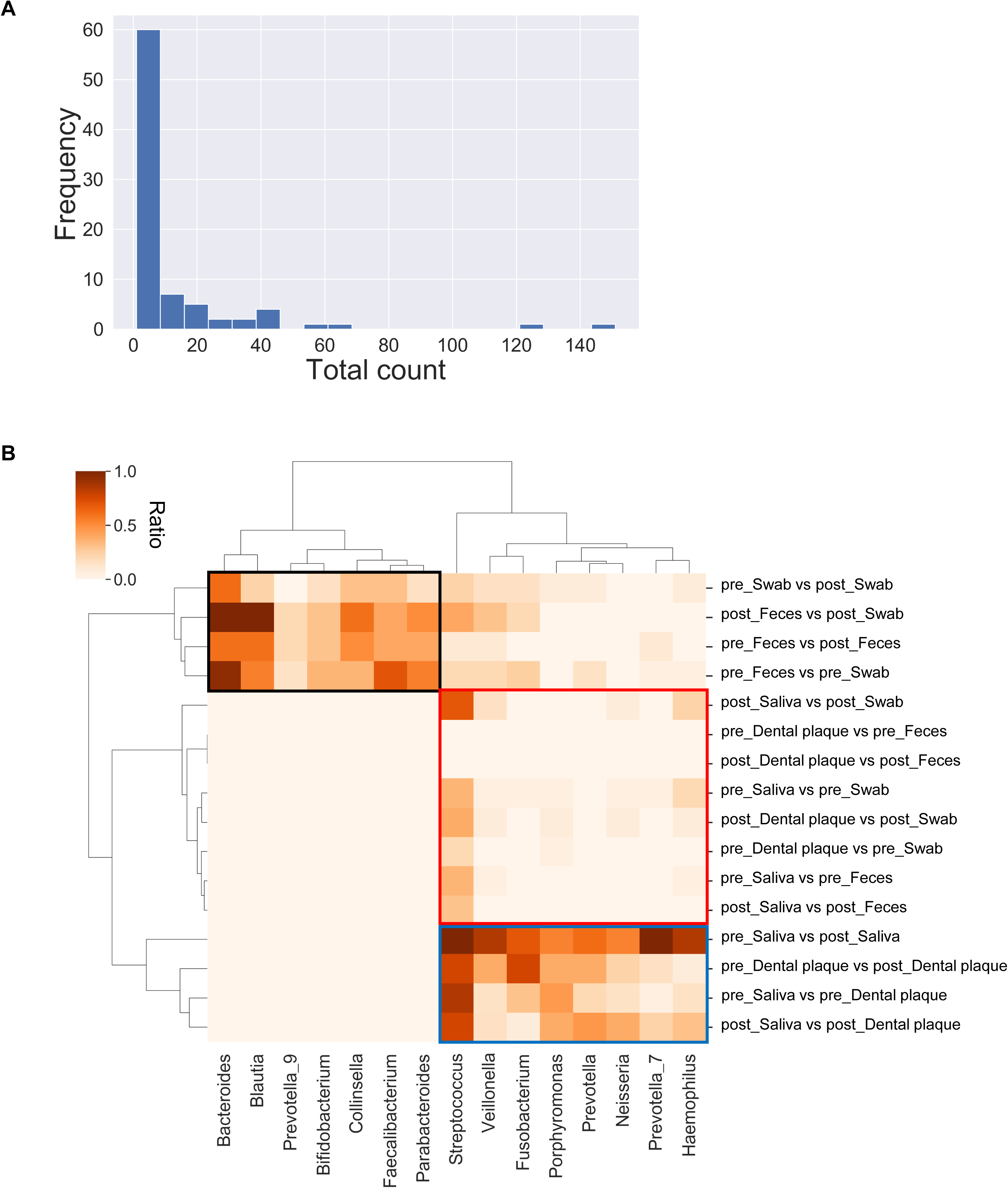
Core microbiome in comparison between each sample. (A) Distribution of the number of patients sharing a genus of bacteria with relative abundance (>1%) in comparison between each sample. (B) Heatmap of the percentage of patients sharing a genus of bacteria with relative abundance (>1%) in comparison between each sample. Top 15 genera were extracted in a number of patients sharing the genus. The horizontal axis is the genus, while the vertical axis is the comparison between each sample.

Hierarchical clustering of the ratios of the top 15 genera revealed relationships between each comparison and the core microbiome (**Fig.3B**, **Supplementary Table 3**). Clustering divided each comparison into intra-intestinal environment, comparisons between intestinal and oral environments, and intra-oral environment (**Fig. 3B**). Genera with high ratios for intestinal comparisons were distinct from others. The core microbiome exhibited a high ratio within the same environment, such as *Bacteroides* and *Blautia* for intra-intestinal comparisons (**Fig. 3B** black square), and *Streptococcus* and *Porphyromonas* for intra-oral comparisons (**Fig. 3B** blue square). Interestingly, a ratio above zero was observed for the core microbiome between different environments, such as *Veillonella* and *Porphyromonas* (**Fig. 3B** red square), indicating potential translocation of bacteria between oral and intestinal environments.

### Shared ASVs analysis suggested the translocation of periodontal pathogens to the gut

To investigate microbial translocations between oral and intestinal environments, shared amplicon sequence variant (ASV) analysis was done to detect intra-individual sharing at the ASV level. Our analysis revealed that many patients shared ASVs of *Streptococcus* between oral and intestinal environments pre– and post-operation (**Supplementary** Fig. 2, **Supplementary Tables 4**,**5**). Additionally, *Haemophilus* and periodontal pathogens including *Porphyromonas*, *Prevotella_7*, *Fusobacterium*, *Neisseria*, and *Veillonella* were detected between saliva and subgingival dental plaques before and after the operation (**Fig. 4**, **Tables S4**,**5**). Specifically, preoperatively, *Haemophilus* and periodontal pathogens, including *Fusobacterium*, *Neisseria*, *Porphyromonas*, *Prevotella_7*, and *Veillonella*, were detected in saliva and swab samples, but postoperatively, *Fusobacterium*, *Porphyromonas*, and *Prevotella_7* were not detected under the threshold. In subject 9, 14, and 16, two types of ASVs of *Veillonella* (ID: Ve_13 and Ve_16) were detected in samples both pre– and postoperatively. *Haemophilus* and *Veillonella* were also detected between saliva and feces before the operation. Furthermore, the periodontal pathogen *Porphyromonas* was shared between subgingival dental plaque and swab samples both pre– and postoperatively. Postoperatively, *Haemophilus*, *Neisseria*, and *Veillonella* were also shared between subgingival dental plaque and swab samples. These findings collectively suggest that oral bacteria can translocate to the gut, although the occurrence rate of periodontal pathogens is lower compared to *Streptococcus*.

**Figure 4.**
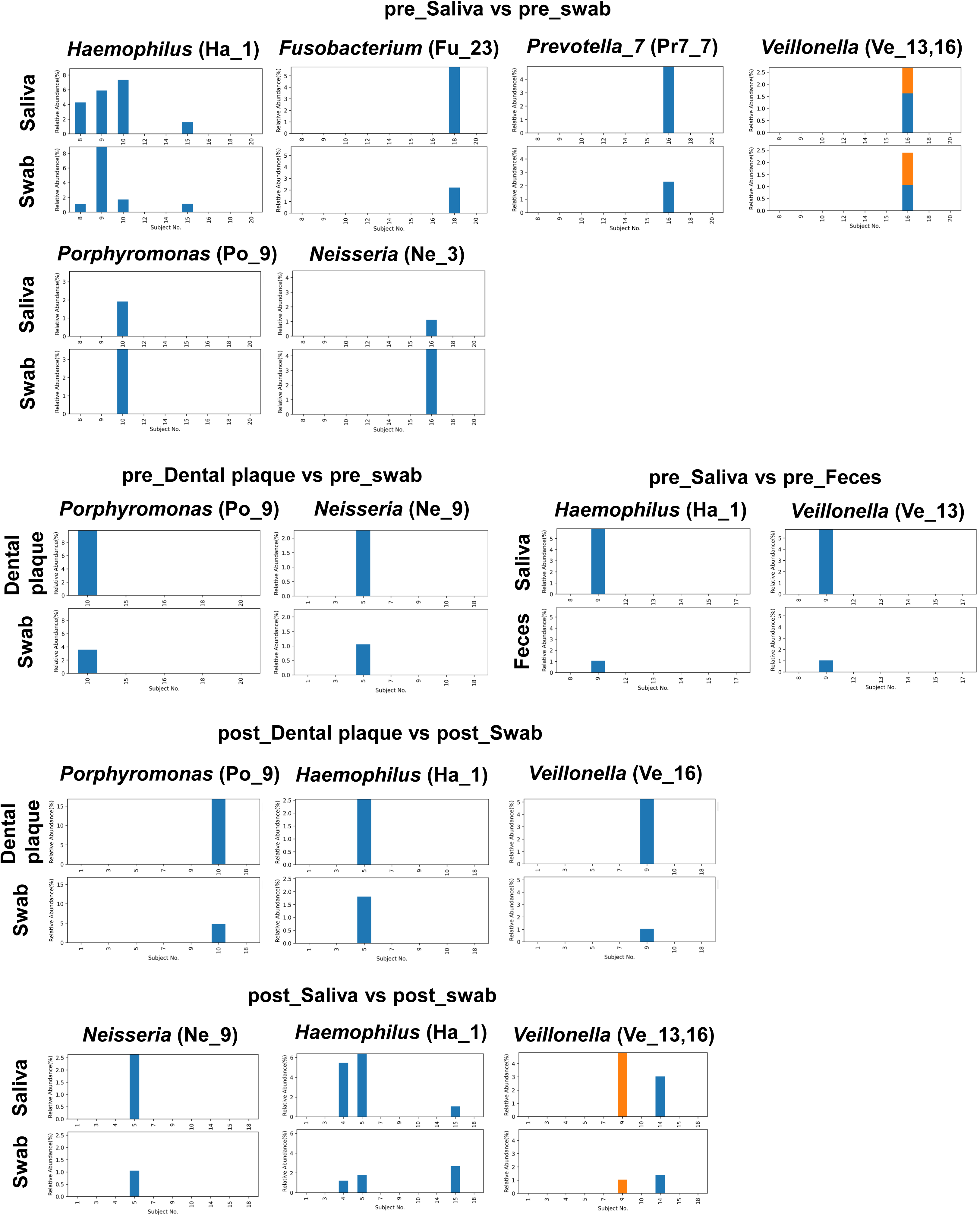
Shared ASVs in comparison between each sample. The bar plot indicates the relative abundance of ASVs in comparison between each sample. The color indicates each ASV. The label name of legend is described in Supplementary Table 4.

## Discussion

In this study, oral and intestinal microbiota from pre– and postoperative periods of endoscopic colorectal polypectomy were analyzed using several bioinformatic approaches. Core microbiome and shared ASV analyses revealed bacterial sequence sharing between oral and intestinal environments, suggesting bacterial transmission from the oral cavity to the gut in Japanese patients with colorectal polyps.

As depicted in the bacterial composition of each sample (**Fig. 1**), *Fusobacterium* and *Veillonella* at the genus level were observed in feces and swab of intestinal tissue. Both *Fusobacterium* and *Veillonella* are commensal bacteria in both the oral and gut microbiota and are implicated in periodontal disease^30,31^. *Fusobacterium*, an anaerobic gram-negative bacterium, is detected in the gastrointestinal tract, including the stomach, colon, and ileal pouch^32^. *Fusobacterium nucleatum* is significant in subgingival dental plaque and associated with periodontitis and CRC^33^. *Veillonella parvula* is present in the oral cavity and has been found to colonize the intestine, particularly in patients with inflammatory bowel disease^34^. These findings suggest that both *Fusobacterium* and *Veillonella* translocate from the oral cavity to the gut, potentially contributing to gastrointestinal tract disorders.

Regarding beta diversity analysis, oral samples (saliva and subgingival dental plaque) and intestinal samples (feces and swab of intestinal tissue) were segregated as expected.^14^ However, no significant differences were observed between feces (regarded as LAM) and swab (regarded as MAM) samples (**Fig. 2B**). It is commonly observed that LAM and MAM exhibit differences in their composition^14,35,36^. For instance, Miyauchi et al.^14^ conducted a comparative analysis between LAM derived from feces and MAM samples collected by swabbing the mucosa in healthy volunteers. A weighted UniFrac distance differed between LAM and MAM, and PCoA clearly separated LAM from MAM. Given the biogeographic adjacency of MAM to the intestinal epithelium, MAM provides more precise information in predicting bacteria associated with intestinal diseases. Although it is unclear why no differences were observed between LAM and MAM in this study, technical procedures of sample collection, as well as the process of sample analysis, might influence the results.

Core microbiome analysis is a bioinformatics approach that identifies consistent microbial characteristics across particular populations or environments. In this study, we conducted core microbiome analysis to identify intra-individual sharing of microbial taxa present at the genus level. The results exhibited a ratio above zero between oral and intestinal environments, such as *Streptococcus*, *Veillonella*, *Fusobacterium*, *Porphyromonas*, *Neisseria*, *Prevotella_7*, and *Haemophilus* (**Fig. 3B**, red square), indicating possible translocation of these bacteria between oral and intestinal environments. These results align with previous studies. Schmidt et al.^9^ conducted bioinformatic analysis using a dataset of 753 public and 182 newly sequenced saliva and stool metagenomes from 470 individuals. They profiled 310 prevalent species and concluded that strains of *Streptococcus*, *Veillonella*, *Actinomyces*, and *Haemophilus* were subject to frequent oral-fecal microbial transmission. Additionally, Kageyama et al.^10^ examined 144 pairs of saliva and stool samples collected from community-dwelling adults. Their bioinformatic analysis revealed that *Streptococcus* and *Prevotella* were predominant in both saliva and feces, with *Veillonella* and *Megasphaera* overlapping slightly. The higher oral-gut transmission of *Streptococcus* compared to periodontopathogenic bacteria has also been reported in these studies due to the abundant prevalence of *Streptococcus* in the oral cavity; the number of bacteria transmitted to the gut would be proportional to the prevalence of the bacteria at the origin site.

In this study, shared ASV analysis was done to predict bacteria that translocated from oral to gut. This analysis method is frequently used as an alternative to operational taxonomic units for characterizing and comparing microbial communities^26,27^. The shared ASV analysis demonstrated that some periodontal pathogens, such as *Porphyromonas*, *Haemophilus*, *Neisseria*, and *Veillonella*, were shared between oral and intestinal samples (**Fig. 4**), implying the translocation of periodontal pathogens to the gut. However, the ASV analysis using the V3-V4 region of 16S ribosomal RNA in this study includes uncertainty regarding taxonomic discrimination; therefore, long-read sequencing of the full-length 16S ribosomal RNA gene could provide more reliable results with higher resolution.

Another finding of the ASV analysis in this study is that the pre– and postoperative comparisons showed no characteristic microbiota signature, consistent with previous reports by Yu et al^37^. They compared alterations in microbial diversity and structure of stool samples before endoscopic therapy and approximately 3 months after resection, respectively. Adenoma excision brought no significant alterations in microbial composition and diversity. Furthermore, Sze et al.^38^ compared fecal microbiota of patients before and after treatment for colonic lesions and found no significant change in diversity in the pre– and post-treatment samples. However, the intestinal existence of *F. nucleatum*, acknowledged as a biomarker for CRC, has been characterized by multiple studies. There is notable prevalence of this species in adenoma tissues and exhibits a gradual increase in abundance with the progression of colorectal carcinogenesis^39,40^.

Despite our efforts to provide a comprehensive understanding of the potential transmission of oral bacteria to the gut in Japanese patients with colorectal polyps, several limitations remain to be addressed in future studies. First, we were not able to enroll a large number of participants. Furthermore, some postoperative samples could not be collected due to unexpected hospital transfers and/or changes in treatment plans of patients, hence the need to perform analysis on a larger number of samples in the future. Second, our microbial analysis employed the V3–V4 region of 16S ribosomal RNA, which includes uncertainty regarding taxonomic discrimination; therefore, a combination of long-read sequencing and shallow metagenomics could provide more information at the species level. Additionally, employing metagenomic analysis for functional characterization may advance our understanding of microbial involvement in CRC pathogenesis. Finally, this is only an observational study, and an interventional study is needed to obtain a more definitive understanding of the causal relationship between periodontal pathogens and CRC.

## Conclusions

The core microbiome and shared ASV analyses showed that several genes are shared between oral and intestinal environments in patients with colorectal polyps, indicating the oral-gut translocation of periodontitis-related bacterium. Further large-scale studies are needed to elucidate its involvement in CRC.

## Declarations

### Ethics approval and consent to participate

This study was conducted according to the principles of the Declaration of Helsinki and was approved by the Ethics Committee of Niigata University Medical and Dental Hospital (Approval No: 2021-0018 and G2022-0001).

### Consent for publication

Informed consent was obtained from all the subjects involved in the study. Written informed consent was obtained from the patients for publication of this paper.

### Availability of data and materials

The datasets generated through 16S rRNA amplicon sequencing are available and deposited in the NCBI Sequence Read Archive database under accession numbers: DRR542882-DRR543010 and BioProject: PRJDB17852

## Authors’ contributions

NT, KS and HS designed and set up the experiments with the support of HM and KT. NT, MY, and KS wrote the manuscript with the support of ST and KT. KI, YI, and MY performed microbiome analysis and statistical analyses. NT, KS, TT, SM, and YN collected the oral samples. NN and KT collected the intestinal samples. All authors have contributed to the manuscript and approved the submitted version.

## Acknowledgements

All authors thank Morgenrot Inc. for providing the computational environment for the analysis. We would like to thank Dr. Suguru Fujita at The University of Tokyo for giving insightful comments. The authors would like to thank Enago (www.enago.jp) for the English language review.

## Supporting Material

**Supplementary Table 1**. Number of patients for each comparison.

**Supplementary Table 2**. Ratio of patients sharing a genus of bacteria with a relative abundance (>1%) in all comparison between each sample.

**Supplementary Table 3**. Ratio of patients sharing a genus of bacteria with a relative abundance (>1%) in comparison between each sample.

**Supplementary Table 4**. ASV ID and alternative ID.

**Supplementary Table 5**. Shared amplicon sequence variant (ASV) analysis in comparison between each sample.

## Figure legends

**Supplementary Figure 1.**
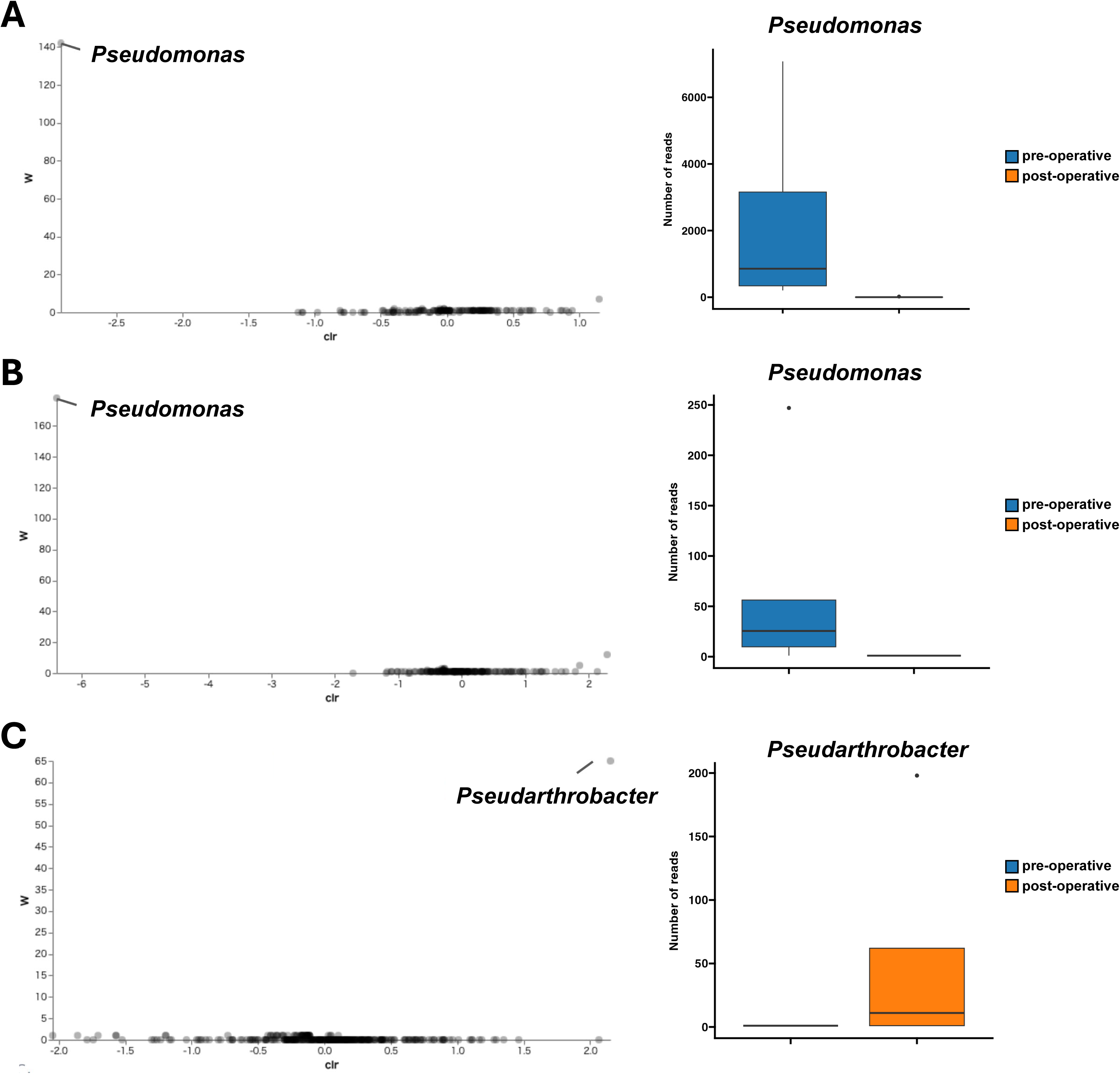
Differentially abundant microbial taxa identified by Analysis of Composition of Microbiomes (ANCOM). Volcano plots of the difference in abundance in each sample, comparing patients before and after surgery. Clr (x-axis) is the measure of the effect size difference for a particular species between the two conditions; W-statistic (y-axis) is the strength of the ANCOM test for the number of species tested. Boxplots are pre– and post-surgery abundances for the genus with the highest W-statistic in each sample (A: Saliva, B: Dental plaque, C: Swab of intestinal tissue).

**Supplementary Figure 2.**
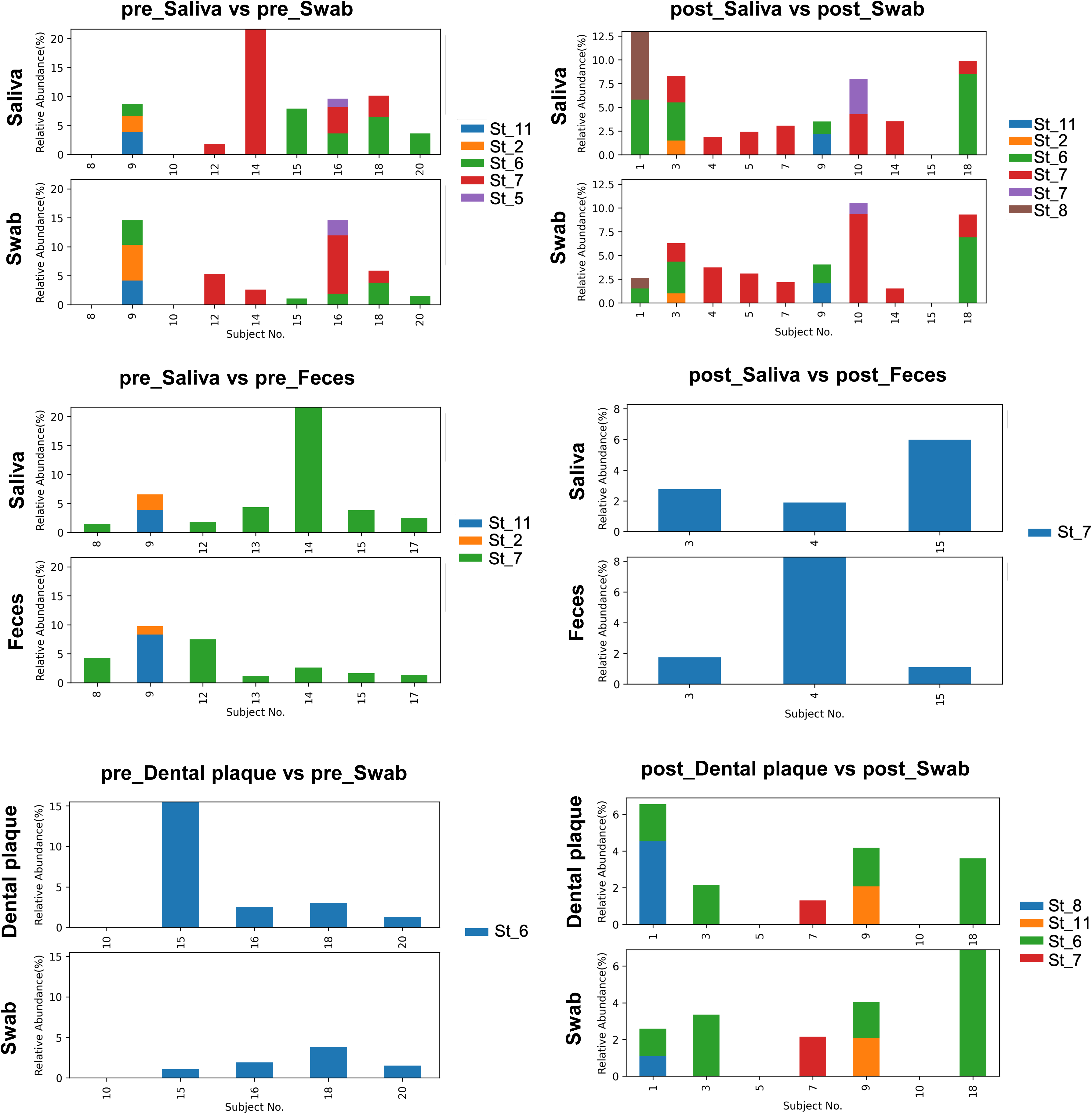
Shared ASVs in comparison between each sample for Streptococcus. The bar plot indicates the relative abundance of ASVs in comparison between each sample for Streptococcus. The color indicates each asv. The label name of legend is described in Supplementary Table 4.

